# Mining the archives: a cross-platform analysis of gene expression profiles in archival formalin-fixed paraffin-embedded (FFPE) tissue

**DOI:** 10.1101/019794

**Authors:** Anna Francina Webster, Paul Zumbo, Jennifer Fostel, Jorge Gandara, Susan D. Hester, Leslie Recio, Andrew Williams, Chales E. Wood, Carole L. Yauk, Christopher E. Mason

## Abstract

Formalin-fixed paraffin-embedded (FFPE) tissue samples represent a potentially invaluable resource for transcriptomic-based research into the molecular basis of disease. However, use of FFPE samples in gene expression studies has been limited by technical challenges resulting from degradation of nucleic acids. Here we evaluated gene expression profiles derived from fresh-frozen (FRO) and FFPE mouse liver tissues using two DNA microarray protocols and two whole transcriptome sequencing (RNA-seq) library preparation methodologies. The ribo-depletion protocol outperformed the other three methods by having the highest correlations of differentially expressed genes (DEGs) and best overlap of pathways between FRO and FFPE groups. We next tested the effect of sample time in formalin (18 hours or 3 weeks) on gene expression profiles. Hierarchical clustering of the datasets indicated that test article treatment, and not preservation method, was the main driver of gene expression profiles. Meta- and pathway analyses indicated that biological responses were generally consistent for 18-hour and 3-week FFPE samples compared to FRO samples. However, clear erosion of signal intensity with time in formalin was evident, and DEG numbers differed by platform and preservation method. Lastly, we investigated the effect of age in FFPE block on genomic profiles. RNA-seq analysis of 8-, 19-, and 26-year-old control blocks using the ribo-depletion protocol resulted in comparable quality metrics, including expected distributions of mapped reads to exonic, UTR, intronic, and ribosomal fractions of the transcriptome. Overall, our results suggest that FFPE samples are appropriate for use in genomic studies in which frozen samples are not available, and that ribo-depletion RNA-seq is the preferred method for this type of analysis in archival and long-aged FFPE samples.

**Author contributions:** 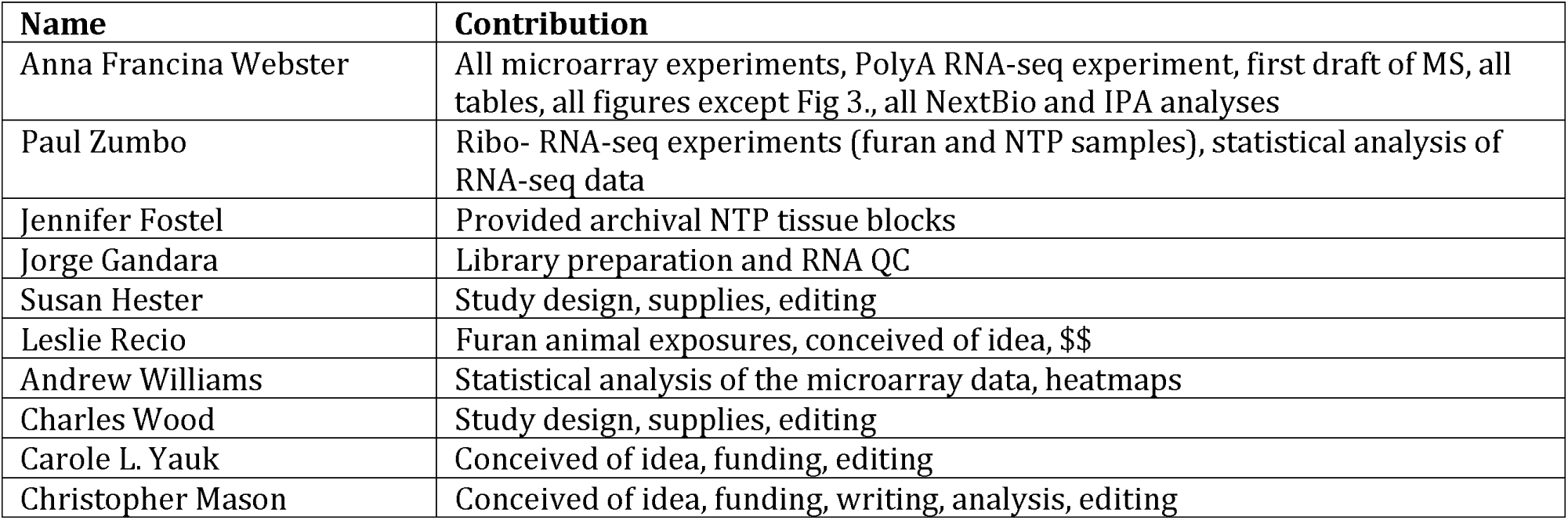

## Introduction

Tissue repositories have immense untapped potential in translational research. Applications of archival tissue resources range from biomarker discovery and toxicogenomic profiling to large-scale studies of genome-disease interactions. Curated sample banks house diverse tissue types from different models or study populations often linked with detailed clinical or pathologic outcomes. In many cases tissue archives may contain unique samples from animal bioassays, clinical trials, or epidemiologic studies that may be different or impossible to repeat. Despite this promise, direct use of archival samples for transcriptomic profiling has been relatively limited to date. The majority of archival tissues are stored in formalin-fixed paraffin-embedded (FFPE) blocks, which preserve tissue architecture for histopathological analysis and allow for tissue storage at room-temperature. However, formalin treatment degrades RNA (through cross-linking and fragmentation), which significantly impairs molecular analyses (Bass et al. 2014; Klopfleisch et al. 2011; Farragher et al. 2008). These RNA effects can result in inconsistent genomic data and present important technical and analytical challenges when working with FFPE samples. Nevertheless, studies have demonstrated the value of using microarrays to analyze FFPE samples for understanding the molecular basis of a variety of cancers (Sadi et al. 2011; Jacobson et al. 2011; Budczies et al. 2011; Linton et al. 2012). As technologies continue to improve, so too does the prospect of FFPE genomics.

Recently, whole-transcriptome RNA sequencing (RNA-seq) methodologies have been developed as a more precise and less biased measurement of transcript levels that may overcome issues associated with highly fragmented or degraded RNA. Compared to conventional DNA microarrays, RNA-seq enriches for many additional fragments as it is not restricted to predefined probes and has (in principle) no limitations to dynamic range (Li et al. 2014b; SEQC/MAQC-III Consortium 2014). Previous work has shown concordance between gene expression profiles produced by microarrays and by RNA-seq for fresh-frozen (FRO) and FFPE samples (Marioni et al. 2008; Malone and Oliver 2011; Wang et al. 2014). Recent studies investigating the use of RNA-seq in FFPE samples have also shown promising results (Hedegaard et al. 2014; Auerbach et al. 2014; Zhao et al. 2014a). To extend these findings, the HESI Committee on Application of Genomics to Mechanism-Based Risk Assessment coordinated a working group to assess the utility of various gene expression profiling methods for FFPE tissues. The main goals of this group were to evaluate current RNA-seq and microarray methods in paired FRO and FFPE samples, and investigate different preanalytical and experimental factors influencing FFPE results.

The reference compound for this study was furan, which is a known liver carcinogen in rodents (NTP 1993; Moser et al. 2009; IARC 1995). Typically, humans are exposed to furan by consuming heat-treated foods or inhaling combustion emissions (Moro et al. 2012a; IARC 1995). Furan is metabolised by cytochrome P450 2E1 (CYP2E1) into an electrophilic and cytotoxic metabolite, cis-2-butene-1,4-dial (BDA) (Kedderis et al. 1993; Kedderis and Held 1996). The carcinogenicity of furan has been attributed to BDA-induced cytotoxicity followed by dysregulated regenerative proliferation in the liver (Fransson-Steen et al. 1997; Moser et al. 2009). We recently confirmed this mode of action using DNA microarray analysis of FRO livers taken from mice that were sub-chronically exposed to both carcinogenic and non-carcinogenic doses of furan alongside solvent-exposed controls (Jackson et al. 2014). We proposed that Cyp2E1 protein stabilization and metabolism of furan lead to chronic activation of the Nrf2 Oxidative Stress Response pathway, which is a primary driver in furan-induced carcinogenic transformation. Here we extend this study by investigating whether the same furan-dependent molecular changes can be detected in paired FFPE samples.

In order to evaluate genomics methods for FFPE tissues, we compared gene expression profiles from paired FRO and FFPE-preserved liver tissues from female B6C3F1 mice that had been exposed for three weeks to a carcinogenic dose of furan. Gene expression profiling was completed using four protocols: Illumina Hi-seq RNA-seq (ribo-depletion and polyA-enrichment formats) and Agilent SurePrint G3 Mouse GE 8x60K microarrays (one-and two-color formats). Next, we investigated the effect of tissue time-in-formalin on the quality of transcriptomic data given that different institutions have different time-in-formalin protocols. For this aim, we used furan-exposed paired mouse liver samples that had been preserved in formalin for either 18 hours or 3 weeks. Finally, to better understand the effects of time-in-paraffin, we used the RNA-seq ribo-depletion protocol to evaluate rat liver samples that were obtained from the National Toxicology Program (NTP) archives and had been in storage for up to 26 years. Our results provide a comprehensive comparison of genomics methodologies for FFPE samples and recommendations for the most appropriate approach for the study of archival specimens.

## Results

### Gene expression profiles

Our initial aim was to evaluate transcriptomic profiles obtained using different protocols. The mean RNA integrity numbers (RIN) for the FRO, FFPE-18hr, and FFPE-3wk samples were 9.3 (range: 8.9-9.5), 2.1 (range: 1.9-2.3), and 2.2 (range: 1.3-2.5), respectively. RNA integrity values are reported in **Suppl. Table S1.** In order to make fair comparisons between RNA-seq and microarray datasets, RNA-seq reads were only considered if they could be mapped back to a transcript corresponding to a probe on the Agilent SurePrint G3 Mouse GE 8x60K microarray. Mapping produced a list of 20,167 RefSeq genes. This list was also used to retrospectively filter the list of microarray-derived differentially expressed genes (DEGs). The numbers of DEGs in the full and mapped gene lists are listed in **Table 1.** With respect to the FRO groups, more DEGs were detected in the FFPE-18hr groups, and fewer were detected in the FFPE-3wk in all experiments apart from the poly-A-enrichment RNA-seq. One-color DNA microarrays consistently detected about twice the number of DEGs as two-color arrays across all sample types. Compared to microarray protocols, RNA-seq DEG numbers were intermediate (FRO and FFPE-3wk) or modestly lower (FFPE-18hr) for the ribo-depletion protocol, and either higher (FRO) or lower (FFPE-18hr and FFPE-3wk) for the poly-A protocol. Applying a 2-fold cut-off reduced the number of DEGs by at least 75% across FRO and FFPE groups, limiting the ability to perform pathway analysis in some groups.

**Table 1:**
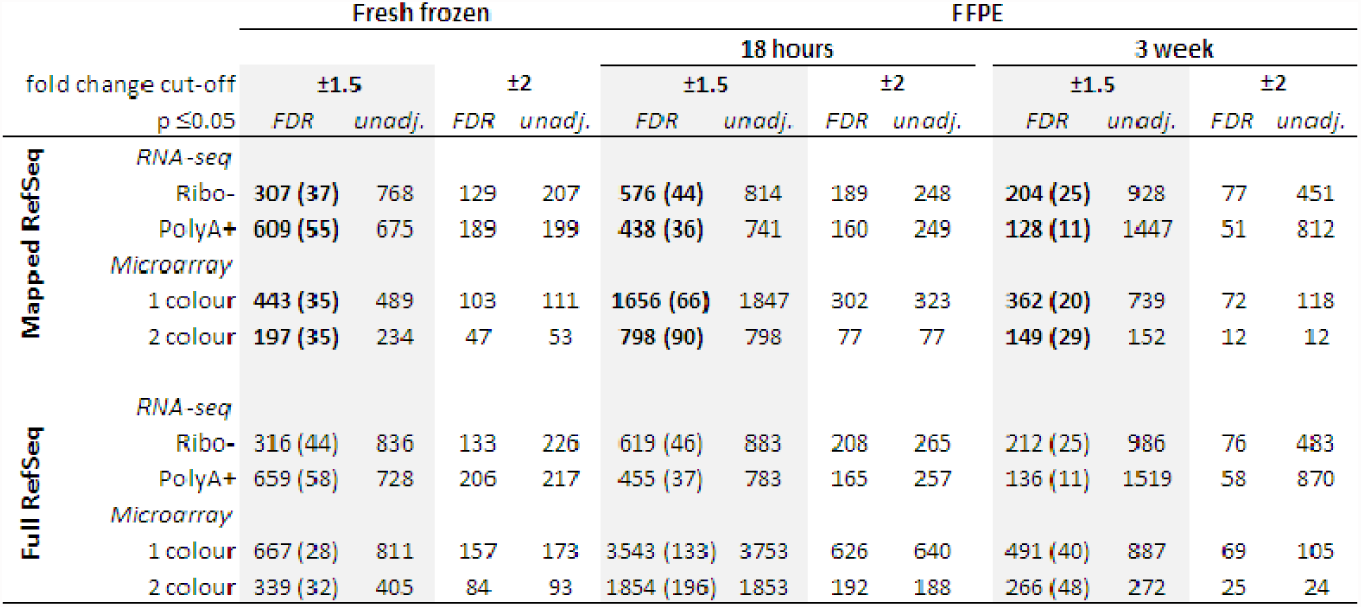
Number of differentially expressed genes. (by unique gene symbol) in each microarray and RNA-seq experiment using different filtering thresholds. The corresponding number of enriched IPA pathways (with p< 0.05 and at least 4 molecules) is indicated in brackets. Lists that were used to derive enriched pathways are indicated by bold text.

Next, we evaluated gene expression correlations by fold-change and sample type (**Fig. 1a**). The R2 correlation coefficients between paired FRO and FFPE samples ranged from 0.29 to 0.93 depending on protocol; all correlations were statistically significant (regression p < 0.05). The correlation between FRO and FFPE-18hr samples was markedly higher for both ribo-depletion and poly-A RNA-seq (but not microarray) protocols compared to correlations between FRO and FFPE-3wk samples, indicating an erosion of signal with time in formalin at the read depths used. Overall, the ribo-depletion method clearly provided the highest correlation between FRO and FFPE samples (at both 18hr and 3wk).

**Figure 1:**
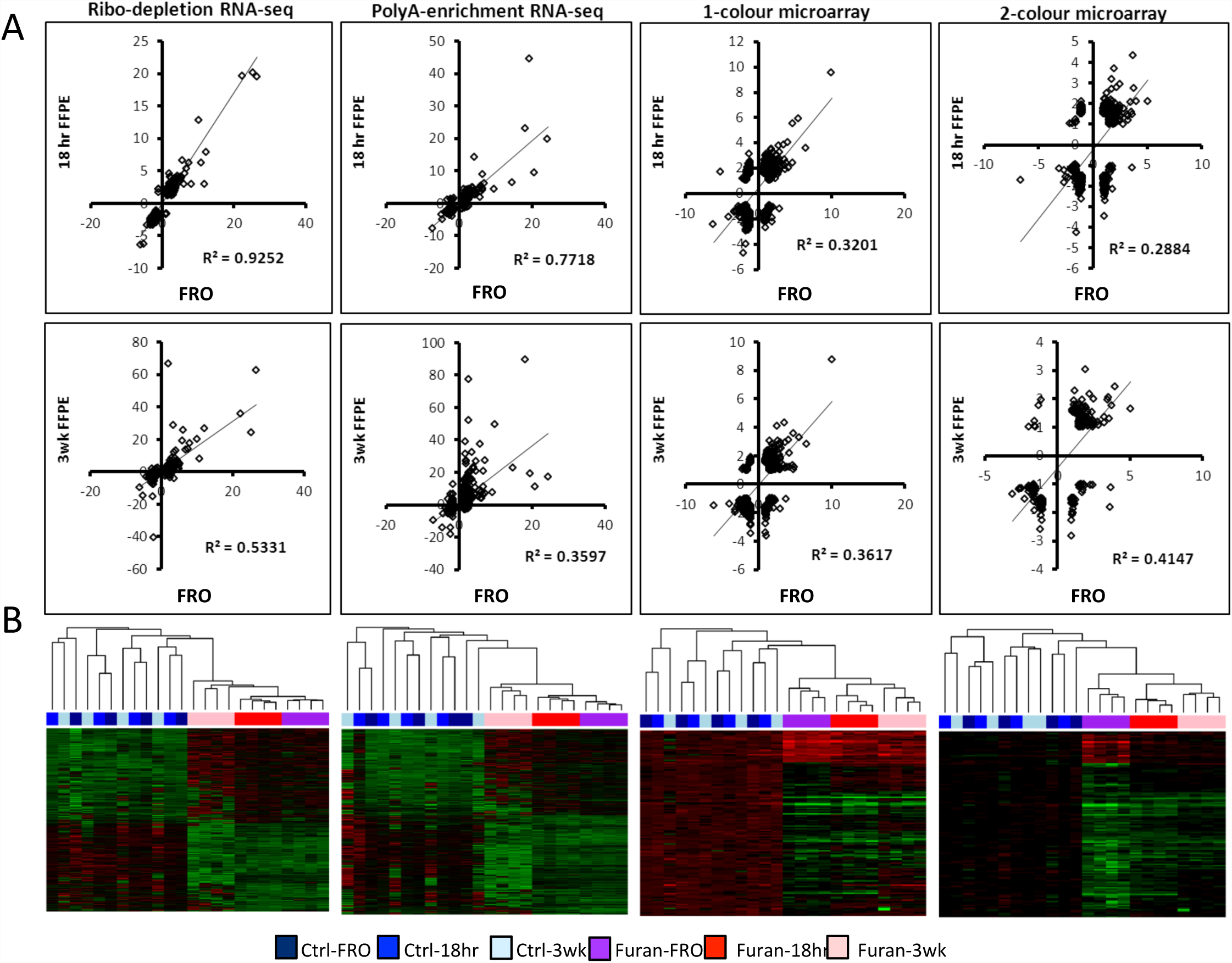
Differential gene expression. Expression profiles following exposure to furan (8 mg/kg/day) in fresh-frozen (FRO), 18 hours in formalin (FFPE-18hr), or 3 weeks in formalin (FFPE-3wk) liver samples evaluated using different DNA microarray and RNA-seq protocols. (A) Correlation analysis of DEG fold changes for FRO *versus* FFPE-18hr (top row) and FRO *versus* FFPE-3wk (bottom row) (p < 0.0001 for all linear regressions). Genes were significantly changed in at least one list (FDR p < 0.05, fold change > ±1.5 compared to control). (B) Hierarchical clustering of all DEGs (FDR p < 0.05, fold change > ±2).

Hierarchical clustering of the datasets demonstrated that furan treatment was the main driver of gene expression profiles (**Fig. 1b**). A decrease in this signal with time-in-formalin is evident from the microarray cluster diagrams, which show a decrease in signal intensity in the furan groups from FRO to FFPE-18hr to FFPE-3wk (the increased amount of black on the heatmaps). Only five genes were up-regulated across all 12 datasets (*Osgin1, Tnfsfr12a, Srxn1, Ephx1,* and *Chka*). Genes up-regulated in 11/12 datasets were: *Cdkn1a, Jun, Dusp6, Hmox1, Brd2, Ugt2b35, Gclc, and Trp53inp1;* and in 10/12 datasets: Tubb2a, Phlda3, Gdf15, Gsta2, Zmat3, Ugdh, Rcan1, Rbp1, Apoc2, and Ces2e. For the four FRO datasets (ribo-, poly-A+, 1-color and 2-color), the overlap of shared DEGs was: 78 (4/4 FRO datasets), 101 (3/4), and 206 (2/4). For the FFPE-18hr groups, these number were: 54 (4/4), 63 (3/4), and 807 (2/4). And, for the FFPE-3wks groups: 7 (4/4), 16 (3/4), and 112 (2/4); demonstrating an erosion of quality with time in formalin. The DEG overlap for the three preservation methods (FRO, FFPE-18hr, and FFPE-3wk) for ribo-depletion was: 86 (3/3) and 165 (2/3); for poly-A-enrichment: 43 (3/3) and 233 (2/3); one-color: 45 (3/3) and 167 (2/3); and two-color: 18 (3/3) and 76 (2/3). Therefore, once again, the ribo-depletion protocol was the most consistent between groups.

### Pathway analysis

Pathway analysis of each DEG list resulted in a large number of enriched pathways (**Table 1**), which were ranked for significance in IPA using the –log(p-value) of a Fisher’s Exact test. To simplify inter-group comparisons, we first considered the top 15 pathways from each group by sample type (FRO, FFPE-18hr, FFPE-3wk). Using FRO as a reference group, the overlap was: 10/15 (ribo-depletion), 5/15 (polyA-enrichment), 4/15 (one-color microarray), 2/15 (two color microarray) (**Fig. 2,** bar charts). Once again, the greatest consistency among sample types was found with the ribo-depletion method. We then compared all enriched pathways between FRO and FFPE for each platform. The percent of FRO pathways that also occurred in FFPE-18hr were: 49% (ribo-depletion), 53% (poly-A-enrichment), 59% (1-color), and 52% (2-color) shared pathways. The percent of FRO pathways that also occurred in FFPE-3wk were: 41% (ribo-depletion), 18% (poly-A-enrichment), 24% (1-color), and 23% (2-color) shared pathways (**Fig. 2, Venn diagrams**). Overall, ribo-depletion and poly-A-enrichment were about equal at 18-hours; however, ribo-depletion was better at 3-weeks. One-color out-performed 2-color microarrays at both time-points.

**Figure 2:**
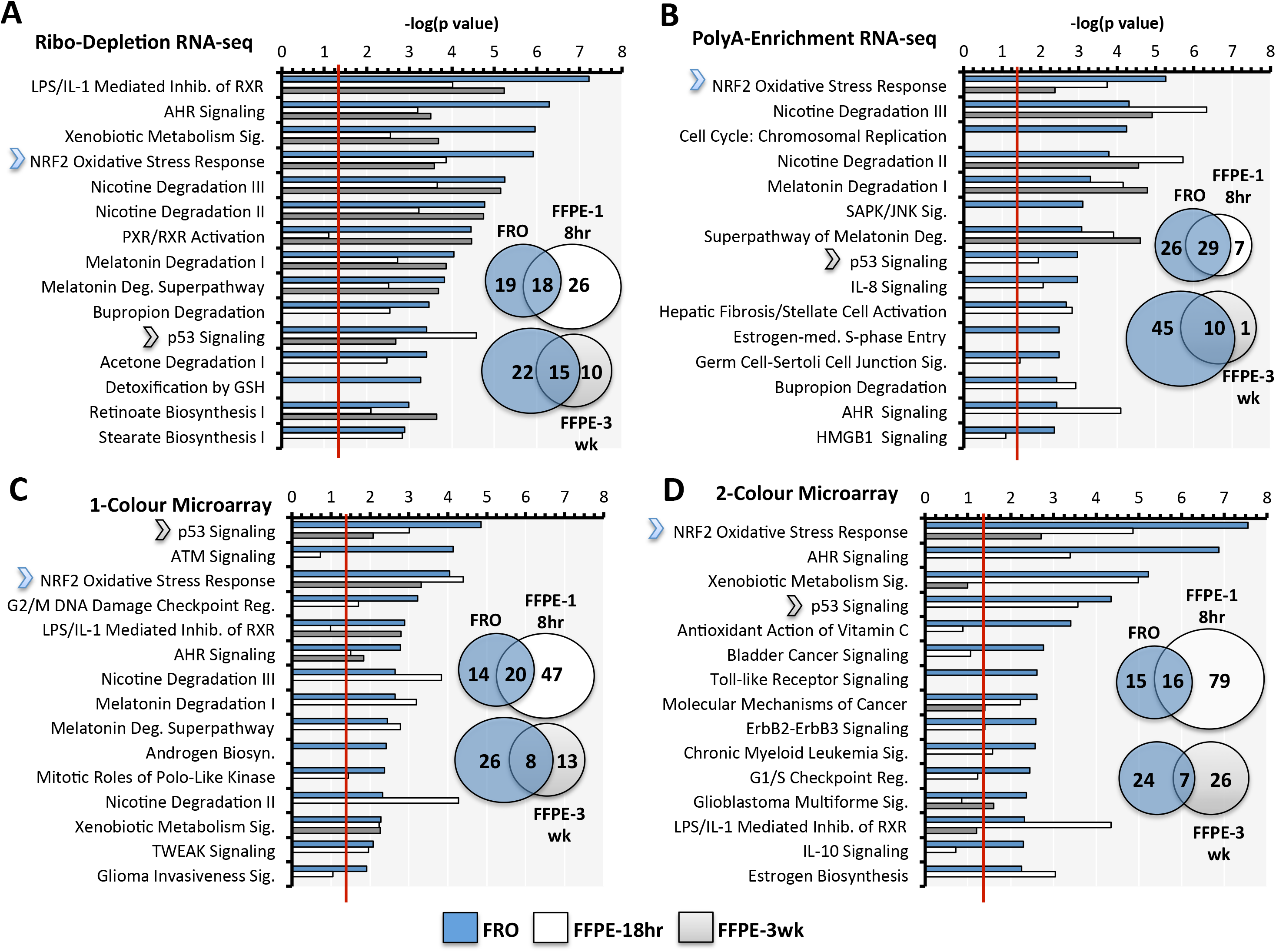
Pathway analysis for (A) ribo-depletion RNA-seq, (B) poly-A-enrichment RNA-seq, (C) one-color microarrays, and (D) two-color microarrays. The top 15 pathways for the fresh-frozen (FRO) groups (blue) are listed vertically. The level of enrichment of the corresponding FFPE groups (white-FFPE-18hrs; grey-FFPE-3wks) is plotted. Pathway significance was calculated in IPA using a Fisher Exact test; −log(p-value) = 1.3 corresponds to a p = 0.05 (indicated in red). Venn diagrams indicate the overlap of each full list. For all analyses, only mapped DEGs with FDR p < 0.05 and fold change > +/-1.5 were used and only pathways with at least 4 DEGs were considered. Nrf2 Oxidative Stress Response was the only pathway that was enriched in all 12 groups (grey stars); p53 was enriched in 11/12 groups (green stars).

Enriched pathways relevant to the mode of action for furan (Jackson et al. 2014) were consistently enriched in FFPE samples (e.g., *Nrf2-mediated Oxidative Stress Response, Xenobiotic Metabolism*, and *p53 Signaling*). Other relevant pathways related to cell cycle (*G2/M DNA damage checkpoint*), cell death (*TWEAK signaling*), inflammatory response (*LPS/IL-1 inhibition of RXR function*), and various cancer pathways were less consistently present across groups. Only the *Nrf2 Oxidative Stress Response* and *p53 Signaling* pathways were shared across all four experimental set-ups, and only the former was enriched in all 12 datasets. The pathway analysis using the unmapped gene lists for each group can be seen in **Suppl. Fig. S1.**

### Meta-analysis of chemicals and diseases produces similar gene expression profiles

We performed a meta-analysis using NextBio to identify chemicals that produce similar changes in gene expression to furan in order to determine if these similarities were detectable in the FFPE furan samples. This meta-analysis was conducted using the data generated by the best-performing protocols from each platform: ribo-depletion RNA-seq and one-color microarray protocols. We performed two chemical meta-analyses. In the first, expression profiles of the top 250 furan-dependent DEGs were compared to those produced in all mouse chemical studies (across the entire NextBio database). Furan was most similar to thioacetamide (on both platforms and across samples), followed by 1-5-napthalenediamine, dimethylnitrosamine, methapyrilene, and diethylnitrosamine (**Table 2**). In the second, we compared our datasets with the changes produced following a 3-week exposure to 26 different chemicals in female B6C3F1 mice by Thomas et al. (2011); this latter study was preselected as a more targeted study subset with the same exposure time, mouse strain, and sex as the current furan study. Compared to the Thomas et al. study, furan produced gene expression changes most similar to malathion, benzofuran, methylene chloride, and 1,5-napthalenediamine. There was a remarkable degree of overlap between FRO and FFPE samples, including identical top five rankings for FRO and FFPE-18hr groups across all NextBio datasets. Overall, the ribo-depletion study produced the best results.

**Table 2:**
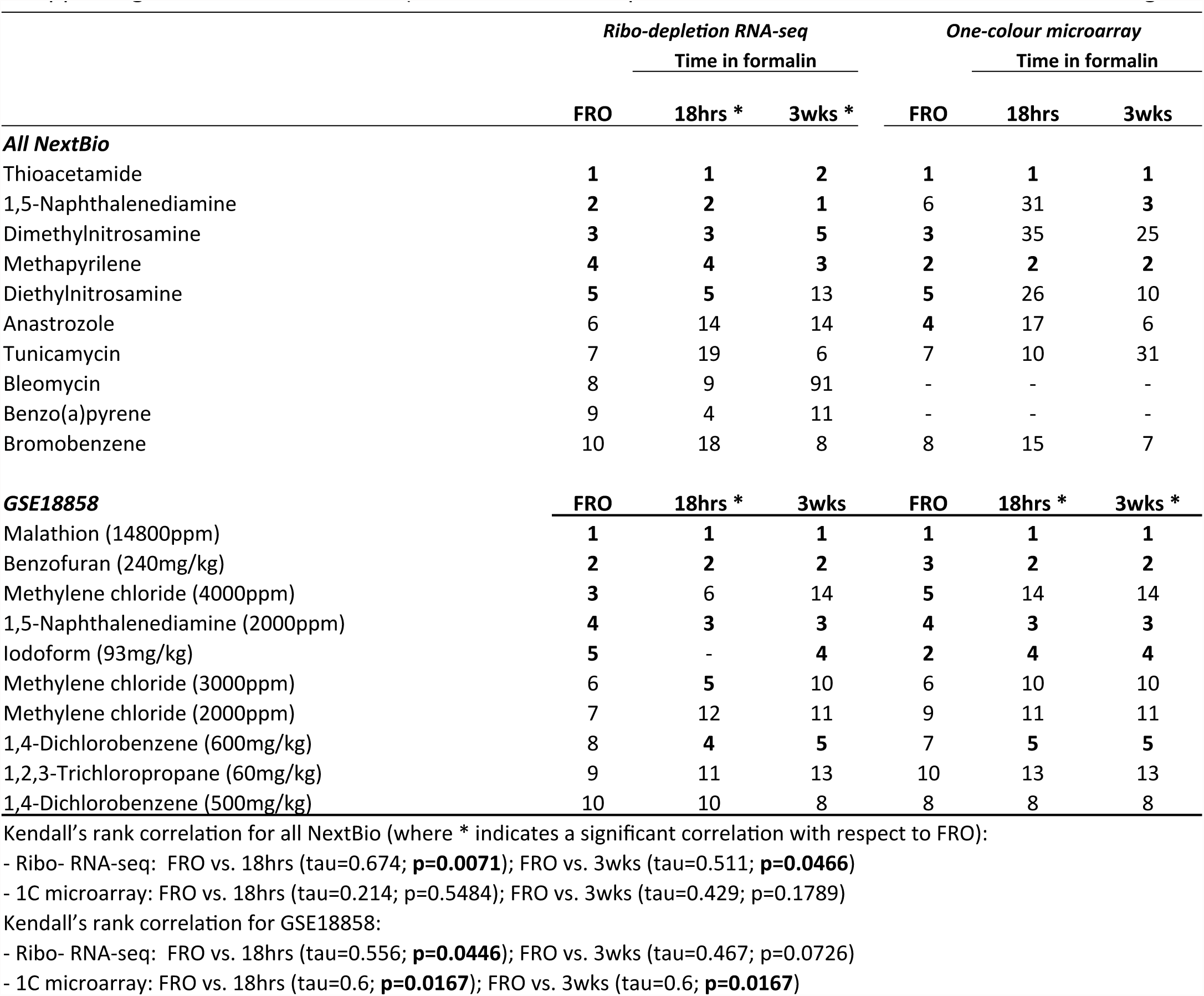
Chemical Signature Analysis. The expression values of the top 250 mapped DEGs (FC > 1.5, FDR p < 0.05 for furan-treated mice *versus* respective controls) stratified by sample group (FRO, FFPE-18hr, FFPE-3wk) were compared to liver gene expression signatures from all NextBio mouse studies (top) and GSE18858 (Thomas et al. 2011; bottom). In the latter, female B6C3F1 mice were exposed to one of 26 chemicals for 13 weeks. The top 5 most correlated datasets for each group are indicated by bold text (where 1 indicates the most correlated study, 2 the second most correlated, etc.); only positively correlated chemicals with at least 3 supporting studies were included (all correlations are p < 0.05, which was calculated in NextBio using a Fisher’s exact test).

A third meta-analysis was performed to identify liver and biliary disease states that produce similar gene expression profiles to furan exposure (**Table 3**). The most correlated studies involved differential gene expression following partial hepatectomy and exposure to carbon tetrachloride (a Cyp2E1 substrate). Both of these are highly consistent with the known mode of action for furan, which includes xenobiotic metabolism by Cyp2E1, cytotoxicity, and regenerative proliferation of the liver. The FRO, FFPE-18hr, and FFPE-3wk groups shared 9/10 (ribo-depletion RNA-seq) or 10/10 (one-color microarray) of the most correlated studies, indicating consistency across the two platforms and three preservation methods for the highest ranking DEGs.

**Table 3:**
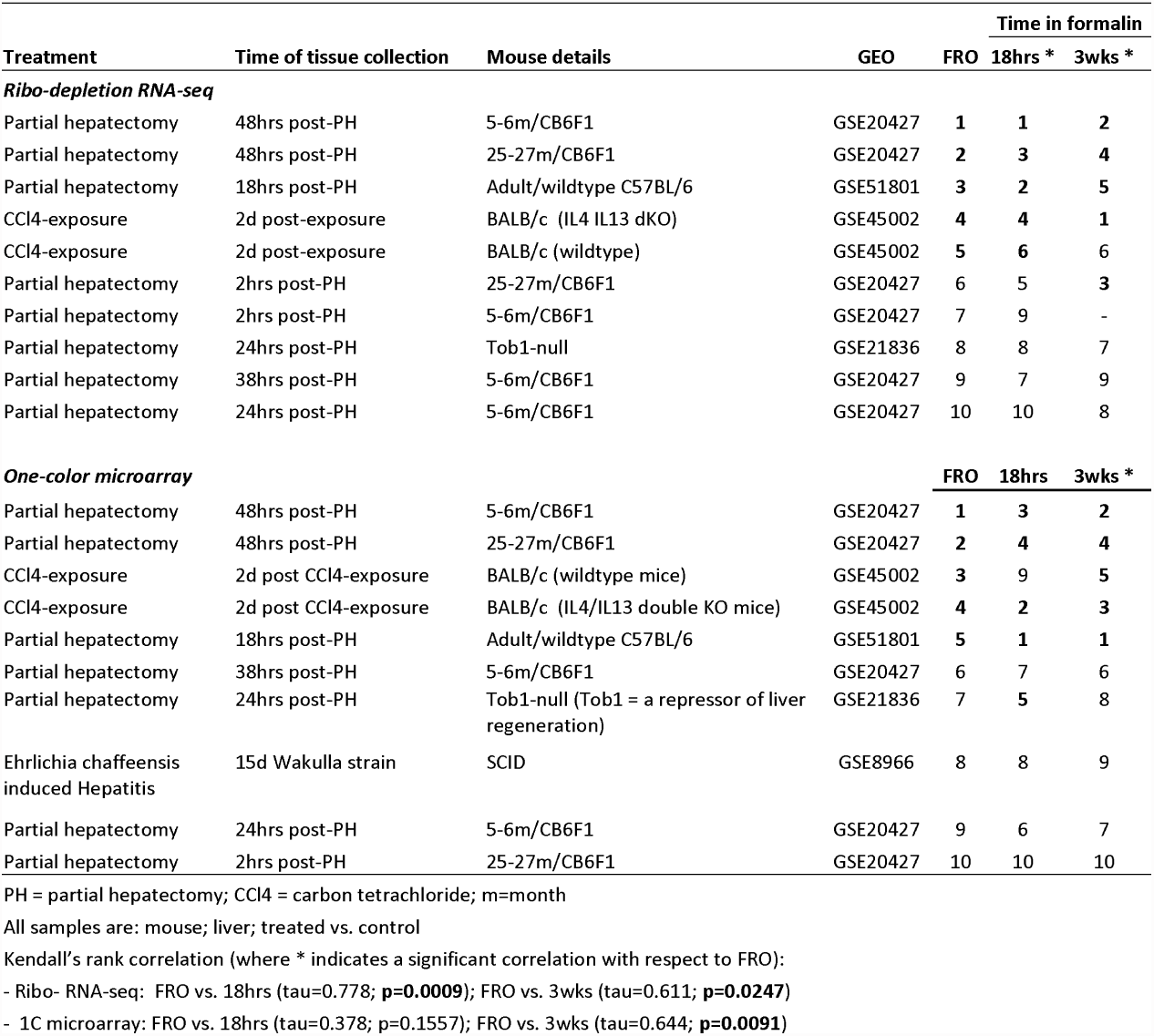
Disease Signature Analysis. The expression values of the top 250 mapped DEGs (FC > 1.5, FDR p < 0.05) in each group (FRO, FFPE-18hr, FFPE-3wk) determined by RNA-seq or microarray analysis were compared to gene expression signatures derived from publically available mouse liver and biliary disease datasets using NextBio. The top 5 most correlated datasets for each group are indicated by bold text (where 1 indicates the most correlated study, 2 the second most correlated, etc.); only positively correlated studies were included. All correlations have a p < 0.05, which was calculated in NextBio using a Fisher’s Exact Test.

### RNA-seq sample profiles

To examine the impact of library preparation on fresh-frozen and FFPE transcriptomic profiles, we prepared libraries using either poly-A-enrichment or ribo-depletion protocols, and sequenced the libraries on an Illumina HiSeq2000 instrument. In order to evaluate the different libraries, we examined different aspects of the RNA-seq data using quality metrics that have been previously developed (Wang et al. 2012, Li et al., 2014), including gene-body coverage uniformity, guanine-cytosine (GC) content distribution, and the distribution of the reads across genic structures (UTRs, exons, introns, or intergenic regions).

We observed that the gene body coverage (calculated as a percentage of reads that covered each nucleotide position of all RefSeq genes scaled to 100 bins) was dependent on both the library preparation method and the sample type (**Fig. 3a**). In particular, because poly-A-enrichment is 3’ biased, and because FFPE samples are more degraded than their fresh-frozen counterparts, the uniformity of gene coverage was biased toward the 3’ end of the gene for poly-A-enriched FFPE libraries. On the other hand, ribo-depleted RNA samples consistently showed more uniform gene coverage than did poly-A-enriched samples; however, after 3 weeks in fixation, even ribo-depleted samples began to show 3’ bias, indicating that 3 weeks in fixation is too long if one is concerned about detecting full-length RNA transcripts.

**Figure 3:**
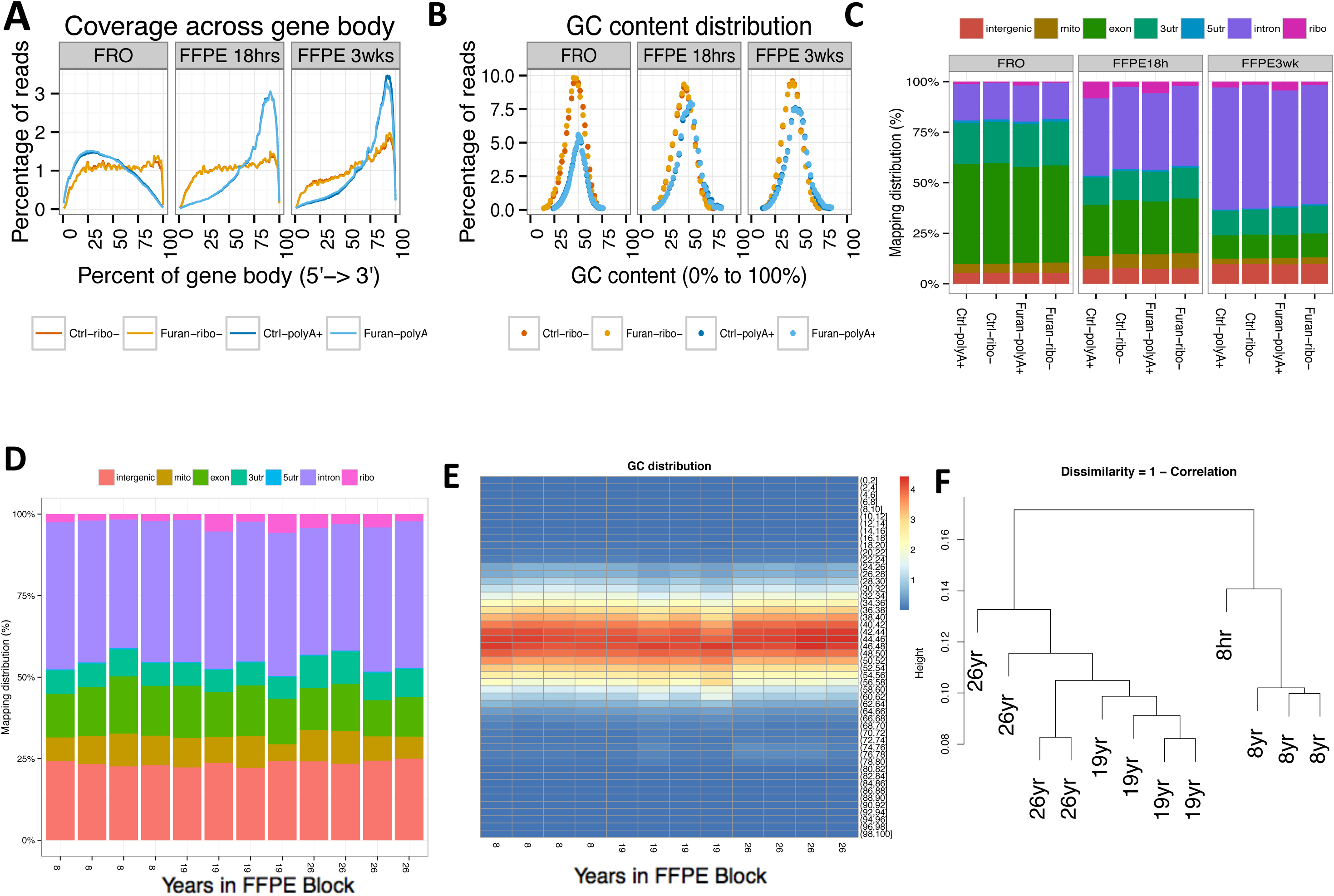
Examination of RNA-seq data quality metrics. (A-C) Mapping distributions and GC content distributions for the furan-treated sampled. (A) The percentage of reads that covers each nucleotide position of all of genes scaled to 100 bins, from 5 ⍰ UTR to 3⍰ UTR for FRO and FFPE samples. The ribo-depleted RNA samples consistently showed more uniform gene coverage than did their poly-A–selected counterparts. (B) GC content distributions for FRO and FFPE samples. (C) The percentage of FRO and FFPE reads that map to various genic regions. Overall, we detect more intronic reads from ribo-depleted RNA samples than from poly-A–enriched libraries; this trend increased as a function of time-in-formalin. (D-E) Mapping distributions and GC content distributions, expressed as a percentage, for 8-, 19-, and 26-year-old control FFPE tissues.

Additionally, we observed that GC content distributions from each sample followed a bell curve, regardless of library preparation protocol (**Fig. 3b**). However, several differences were observed, namely in the widths and heights of those bell curves, likely indicating a difference in the population that was being sampled. Accordingly, the distribution of reads across genic structure (calculated as the fraction of the reads which mapped to either exons, UTRs, introns, or intergenic regions) revealed differences between the two library preparation protocols and sample types (**Fig. 3c**). It is noteworthy that a higher read depth is required for ribo-depletion experiments (compared to poly-A-enrichment) in order to detect equivalent numbers of protein coding genes. In the current experiment the sequencing depth of the ribo-depletion experiment (10-60M reads) was greater than that used in the polyA-enrichment experiment (3.8-11M reads); however, the latter maintained a larger proportion reads mapped to exonic regions (**Fig. 3c**). Overall, ribosomal RNA-depleted libraries consistently showed a larger proportion of introns being sequenced compared to poly-A-enriched samples, as expected (Li *et al.,* 2014). Poly-A-enriched samples, on the other hand, showed increased exon and 3’ UTR sequencing. Compared to FRO samples, poly-A-enriched FFPE samples showed an increasing number of reads mapping to the 3’ UTR as the underlying RNA population degraded (i.e., as a function of time-in-formalin), in line with observed changes in gene body uniformity. FFPE ribo-depleted samples, on the other hand, showed an increasing proportion of reads mapping to introns as a function of fixation time as compared to FRO.

### Long-term archival samples

To examine whether these techniques would be applicable to a wide range of FFPE samples, we tested an additional set of FFPE samples. These archival FFPE samples were obtained from the NTP and had been banked for 8-26 years. It was not possible to obtained archival furan-exposed samples, so the decades-old samples that were tested here represented the control tissue blocks from toxicity studies for beta-picoline (8 years), oxazepam (19 years), and trichlorfon (26 years). These samples were derived from rat; however, it is unlikely that formalin would interact with the genome in a species-specific manner and thus we would expect to see the similar results for mouse. We sequenced each set of samples using the ribo-depletion protocol and compared the expression profiles to standard metrics established by the FDA’s SEQC RNA-seq Consortium (Li et al. 2014a).

We observed that the ribo-depletion protocol was very successful in creating expected distributions of mapped reads to the exonic, UTR, intronic, and ribosomal fractions of the transcriptome (**Fig 3d**). The only aspect of the mapped reads that was high was the percent of mapped reads that were intergenic (violet), which could indicate degradation from the long-banked samples, although we found this trend across all samples. Moreover, the percentage of guanosine-cytosine (GC) from each sample followed a normal distribution (**Fig. 3e**), and did not show any significant shift as a function of treatment or time. Overall, these results demonstrate the potential applicability of these techniques for very old FFPE samples.

## Discussion

Archival FFPE samples provide a unique opportunity to directly link molecular profiles and histopathologic or clinical outcomes without the cost and time associated with new studies. This information could provide more detailed mapping of target pathways, understanding of modes of action, and identification of biomarkers for both environmental chemicals and pharmaceutical agents. However, more accurate and standardized methods for transcriptomic analysis of FFPE tissues are needed. Several recent proof-of-concept studies have evaluated RNA-seq as a novel way to derive gene expression profiles from FFPE or degraded RNA samples and reported promising results (Auerbach et al. 2014; Zhao et al. 2014b; Hedegaard et al. 2014; Spencer et al. 2013; Linton et al. 2012, Li et al, 2014). Our goal here was to expand on these studies by examining specific questions related to the use of RNA-seq in FFPE tissues, including characterizing the performance *versus* microarray platform, discerning the optimal library preparation method, and examining the effects of time-in-formalin and age-in-block. The ribo-depletion RNA-seq method with a relatively brief (18 hour) fixation time provided the strongest correlation between FRO and FFPE tissues at a global transcriptomic level. For highly altered DEGs concordance was also observed across FRO and FFPE sample types for both RNA-seq and microarray platforms. Increased time-in-formalin and age-in-paraffin resulted in loss of signal intensity and other indicators of degradation, but not to the extent that key molecular signatures were lost.

Our first aim was to compare both genomics platforms (two protocols each) in order to develop a pipeline for future FFPE genomics experiments. Across platforms, ribo-depletion RNA-seq provided the strongest gene fold change correlations between FRO and FFPE datasets (**Fig. 1**) and the highest overlap of top pathways between FRO and FFPE groups (**Fig. 2**). Within platforms, ribo-depletion outperformed poly-A-enrichment RNA-seq, and one-color outperformed two-color microarrays. Compared to the poly-A-enrichment protocol, the ribo-depletion protocol produced a more even coverage across all genes and also revealed more low-expressing transcripts. Meta-analyses of the ribo-depletion datasets also showed high concordance in top chemical (**Table 2**) and disease (**Table 3**) signatures. The advantage of the ribo-depletion approach over the poly-A-approach, in the context of FFPE samples, is that it does not rely on the poly-A tail of the mRNA molecules, which is known to be highly modified and degraded by formalin (Farragher et al. 2008; Klopfleisch et al. 2011). The strength of the RNA-seq platform over microarrays is that there is (essentially) an unlimited dynamic range and that reads are not confined by predetermined probe sequences. However, because microarrays have been in use for much longer, the data normalization, processing, and analysis associated with them is more established and straightforward. Collectively, these findings clearly support the ribo-depletion approach for future FFPE genomics experiments.

Our second aim was to examine how formalin fixation impacts RNA quality and the detection of a chemical signature. The process of formalin fixation forms crosslinks between cellular macromolecules and introduces various adducts and other covalent modifications. Sample fixation time varies (from hours to weeks) across laboratories; therefore, an important consideration in planning a retrospective FFPE genomics study is to understand how the initial fixation in formalin will affect the quality of the final gene expression data. Our findings indicate that longer fixation times decrease the RNA yields from paraffin-embedded tissues (in our case 3 weeks in formalin resulted in less than half the RNA yield/section compared to 18 hours in formalin), and increase the RNA input needed during library preparation (see methods). This latter effect is likely due to “non-functional” RNA (e.g. adducts or damaged bases) that could be potentially mitigated by treatments like PreCR or other enzymatic and chemical methods. In addition, an extended time in formalin (3 weeks) led to lower correlation with FRO samples (RNA-seq) (**Fig. 1a**), the detection of fewer DEGs (RNA-seq and microarray) (**Table 1**), and an erosion of signal intensity (**Fig. 1b**) (microarray). It is unclear whether the erosion in signal with time in formalin occurs relatively early in the fixation process, or if signal will continue to decline beyond three weeks in formalin. Given that two-year cancer bioassay samples may be kept in formalin for up to 6 months, it is worth investigating samples held in formalin for longer durations in future experiments.

Samples that stayed in formalin for only 18 hours actually produced a larger number of DEGs than their fresh-frozen counterparts for the ribo-depletion RNA-seq and microarray protocols. Reasons for this higher DEG number are not clear, but this same phenomenon has been observed previously (e.g. Hedegaar et al. (2014), suggesting that these samples have an increased propensity for producing false positive genes. To account for this, it might be worthwhile to perform FFPE genomics experiments in two technical replicates and only treat DEGs that appear in both iterations of the experiment as true positives. While not attempted here, we imagine that this is would be the most practical and useful approach for future studies; particularly microarray studies, which seem to be especially prone to this outcome. Also, RNA-seq is more variable at lower read depths; thus, deeper sequencing is another potential way to increase signal intensity and reduce false positive DEGs in lower-quality FFPE samples. Our results suggest that shorter incubations in formalin result in larger than expected DEG lists, whereas longer incubations in formalin have the opposite effect. Signal intensity appears to decrease with increasing time in formalin. These data indicate that researchers undertaking FFPE genomics studies should, whenever possible, opt for samples that have spent a limited amount of time in formalin. Moreover, our findings highlight the fact that FFPE tissue blocks may vary widely in RNA quality (despite similar RIN profiles), pointing to a need for improved quality metrics specific to FFPE samples.

An important future application of FFPE genomics is the identification of key molecular features in a chemical or drug mode of action. Despite inter-platform and inter-preservation differences in DEG lists, the higher-level biology (pathways and meta-analyses) detected in the FFPE groups in our study (particularly the FFPE-18hr) corresponded well with paired fresh-frozen groups and the expected furan-induced molecular changes. This concordance was strongest for the ribo-depletion RNA-seq protocol. Furan is a well-characterized hepatotoxicant with an extensive toxicological literature, including two cancer bioassays (NTP 1993; Moser et al. 2009) and many xenobiotic metabolism (Burka et al. 1991; Kedderis et al. 1993; Carfagna et al. 1993; Chen et al. 1995; Kedderis and Held 1996; Mugford et al. 1997; Chen et al. 1997; Gates et al. 2012), genotoxicity (Peterson et al. 2000; Byrns et al. 2002; Byrns et al. 2006; Durling et al. 2007; Kellert et al. 2008; Leopardi et al. 2010; Cordelli et al. 2010; Neuwirth et al. 2012; McDaniel et al. 2012; Ding et al. 2012), mode of action (Fransson-Steen et al. 1997; Moser et al. 2009; Hickling et al. 2010; Carthew et al. 2010; Moro et al. 2012b), and exposure/risk (Heppner and Schlatter 2007; Gill et al. 2010; Gill et al. 2011; Moro et al. 2012a) studies, making it an excellent test compound for a retrospective FFPE genomics case study. We recently characterized early molecular key events in the mode of action of furan in female mice using fresh-frozen livers and identified a key role for the *Nrf2 Oxidative Stress Response* pathway (Jackson et al. 2014; Recio et al. 2013). Oxidative stress and damage have been previously reported in response to Cyp2E1 activity and cellular exposure to BDA (furan’s cytotoxic primary metabolite) (Chen et al. 1998; Marí et al. 2001; Cederbaum et al. 2001; Gonzalez 2005; Gong and Cederbaum 2006; Lu and Cederbaum 2008; Hickling et al. 2010). In the current study, the *Nrf2 Oxidative Stress Response* pathway was consistently enriched across all groups and one of the few pathways to be enriched across all platforms. The Nrf2 transcription factor becomes active during periods of cellular oxidative stress (Taguchi et al. 2011), and up-regulates genes involved in cellular redox homeostasis and xenobiotic metabolism (Klaassen and Reisman 2010). Pathways that were enriched less consistently across groups tended to reflect common related processes such as cytochrome p450-mediated metabolism pathways (i.e.: aryl hydrocarbon receptor signaling; degradation pathways for nicotine, melatonin, acetone, burpropion; and biosynthesis pathways for stearate, retinoate, and androgens). Pathways enriched in the FRO groups were typically well-represented in the corresponding FFPE-18hr groups, while the FFPE-3wk groups showed fewer response pathways; however, key events, such as the *Nrf2 Oxidative Stress Response* pathways, were detectable in in both. These findings indicate that samples fixed in formalin for a shorter period of time provide more consistent pathway-level results compared to paired frozen samples.

In addition to pathway analysis, we performed two meta-analyses using NextBio software. This analytic approach showed even more inter-group consistency than the pathway analysis, likely because it does not attempt to break down gene expression profiles into specific events (such as pathways); instead, these meta-analyses make comparisons between entire gene expression datasets that have been generated from different chemical treatments or disease states. The chemical and disease signature meta-analyses are important tools for anchoring data generated from a data-poor compound to the molecular mechanisms found in well-studied compound; for instance, thioacetamide- and carbon tetrachloride-induced gene expression changes were the most similar to those induced by furan, which is not surprising as both compounds are also metabolized in the liver by Cyp2E1 (Manibusan et al. 2007; Kang et al. 2008). Even more compelling is that the most similar disease state was liver regeneration followed by partial hepatectomy. Previous studies have shown that the carcinogenic mode of action for furan includes cytotoxicity followed by dysregulated regenerative proliferation (Moser et al. 2009; Fransson-Steen et al. 1997). These meta-analyses indicate that FFPE samples can provide valuable expression-based information for prioritization of chemicals and next-tier studies.

Lastly, we investigated effects of age-in-paraffin block on RNA quality metrics. No clear effects of age-in-block were observed, supporting the idea that older FFPE samples could be used for transcriptome profiling **(Fig. 3)**. Other researchers have reported RNA-sequencing for 9-year-old and 20-year-old samples (Meng et al. 2013; Hedegaard et al. 2014), but to our knowledge, these are the oldest samples tested to date. While the data from these samples did not have the matching FRO samples for comparison, the data suggest that changes in RNA quality parameters due to age-in-block may be limited. Indeed, decades-old samples appear to be a viable option for FFPE genomics, but additional studies are needed to define optimal preanalytical and bioinformatics approaches.

While there are differences between the molecular profiles of FRO and FFPE samples, our study demonstrates that: (1) RNA-seq with ribo-depletion library preparation is the preferred method for transcriptomic analysis of FFPE tissues (however, one-color microarrays may also be suitable); (2) FFPE samples can be used to obtain reliable transcriptomic information related to a toxicant mode of action (in this case furan); and (3) the RNA profile of a compound is affected by the amount of time that the tissue spends in formalin (although this effect becomes less pronounced with respect to higher-level analyses). Time-in-paraffin did not have a large effect on distributions of mapped reads across control FFPE samples. These findings provide evidence that FFPE samples can be used in genomics studies in absence of existing FRO samples (or when new animal studies are not feasible). **Additional studies could also evaluate whether higher read depth may compensate in part for the loss of signal observed in FFPE samples formalin-fixed for longer periods of time, which could impact work** from archival samples in quantitative applications such as transcriptomic benchmark dose (BMD) estimation. This type of work represents an important opportunity for performing retrospective, phenotypically-anchored research that characterizes chemical- and disease-dependent genomic changes. These chemical/disease signatures can be applied and reapplied (via meta-analysis) to the many data-poor environmental chemicals in need of assessment. This information would also provide a critical bridge between emerging *in vitro* data and health effects established in prior toxicologic and epidemiologic studies, accelerating the input and value of higher-throughput models in safety assessment, biomarker development, and drug discovery.

## Methods

### Frozen and FFPE tissue samples

Frozen and FFPE samples for the microarray/RNA-seq comparison and time-in-formalin analysis were obtained from a recent study of female mice treated with furan (CAS No. 110-00-9) (> 99% pure) (Sigma-Alderich Chemical Co., Milwaukee, WI). Doses of 0 or 8 mg/kg bw furan dissolved in Mazola corn oil were prepared as previously described (Jackson et al. 2014; Recio et al. 2013). Mice were housed, dosed, and sacrificed as previously described (Jackson et al. 2014; Recio et al. 2013). Briefly, 5-6 week-old specific-pathogen-free (SPF) B6C3F1 mice were purchased from Charles River Breeding Laboratories (Portage, ME), and housed in a SPF and Association for Assessment and Accreditation of Laboratory Animal Care (AAALAC)-accredited facility. All experiments were conducted in compliance with the Animal Welfare Act Regulations (9CFR1–4). Mice were handled and treated according to the guidelines provided in the National Institutes of Health (NIH) Guide for the Care and Use of Laboratory Animals (ILAR, 1996; http://dels.nas.edu/ilar/). Mice (n=4 per group) were dosed daily by oral gavage for 21 days. Four hours after the final dose, animals were anesthetized by CO_2_inhalation, and euthanized by exsanguination. Livers were removed and trimmed, and portions were either flash-frozen or fixed in a standard solution of neutral phosphate-buffered formalin. Frozen samples were stored at or below −70°C. Fixed samples were kept in formalin for 18 hours or 3 weeks, and then transferred to 100% ethanol. After ∼72 hours, tissues were processed and then embedded in paraffin using standard histologic procedures. FFPE tissue blocks were stored at room temperature for less than one year before RNA was isolated for this study.

For the time-in-paraffin analysis, FFPE blocks of control liver tissue were obtained from the NTP archive collection. The corresponding slides from each block were re-evaluated on histopathology to confirm that the selected tissue was morphologically normal. Samples were obtained from three different studies in control F344 rats in which tissues had been stored as FFPE blocks for different lengths of time: (1) 8 years (13-week controls for beta-picoline exposure); (2) 19 years (2-year controls for oxazepam exposure); and (3) 26 years (13-week controls for trichlorfon exposure). The difference in study length affects both the age of the animal at sacrifice and also the allowable time in formalin. According to NTP study protocol, for a 13-week study the tissues may only stay in formalin for up to 6 weeks, while for a 2-year study fixed tissues are permitted to remain in formalin for up to 6 months. Recently-collected rat liver was used as FRO control.

### RNA extraction and cDNA synthesis

Total RNA was extracted from FRO tissues using the RNeasy Midi RNA Extraction kit (Qiagen, Mississauga, ON, Canada) as described previously (Jackson et al. 2014; Jackson et al. 2013). Total RNA was extracted from slide-mounted furan and archival FFPE tissues using the FFPE RNeasy kit (Qiagen). One 10 um section was used per FFPE extraction per animal for the 18 hours-in-formalin group. Due to low yields of RNA, two 10 um sections were used per FFPE extraction per animal for the 3 weeks-in-formalin group. RNA extracted from FFPE tissues was assessed for integrity using an Agilent 2100 Bioanalyzer (Agilent Technologies Inc., Mississauga, ON, Canada), quantitated using a NanoDrop Spectrophotometer (Thermo Fisher Scientific Inc., Wilmington, DE, USA), and stored at −80°C (Suppl. Table S1). For cDNA synthesis, RNA (250 ng per sample) was reverse transcribed and amplified using the Whole Transcriptome Amplification kit 2 (WTA2; Sigma, St. Louis, Missouri, USA). cDNA was purified using the QiaQuick PCR clean-up kit (Qiagen).

### Microarray analysis

Microarray analysis for the fresh frozen (FRO) tissues was performed using one-color and two-color protocols (all microarray experiments were conducted in the Yauk laboratory, Ottawa, Canada). The one-color FRO protocol was carried out according to the One-Color Microarray-Based Gene Expression Analysis, Low Input Quick Amp Labeling Protocol, version 6.6 (Agilent Technologies, 2012). Briefly, using the RNA Spike-In Kit, One-Color (Agilent Technologies) and the Low Input Quick Amp Labeling Kit, One-Color (Agilent Technologies), 200 ng of RNA from each sample was used to synthesize, amplify, and Cy3-label cRNA. We labeled 4 samples each for the FRO, 18 hours in formalin FFPE, and 3 weeks in formalin FFPE (control animals 2, 3, 4, 6, and treated animals 41, 42, 43, 44); RNA from animal 3 in the FFPE-3wks group did not label so the number of biological replicates for the FFPE-3wks group was reduced to 3 (we included an additional technical replicate of sample 2 in this group). Labeled cRNA was purified using the RNeasy Mini Kit (Qiagen) and quantified using a NanoDrop Spectrophotometer. Hybridization mixes were prepared using the Gene Expression Hybridization Kit (Agilent Technologies). Cy3-labeled cRNA (600 ng) was hybridized to SurePrint G3 Mouse GE 8 x 60 K microarrays (Agilent Technologies) at 65 °C for 17 hours at 10 rpm.

The two-color FRO protocol was previously described in (Jackson et al. 2014). Briefly, using a reference design, 200 ng of sample RNA and 200 ng of mouse universal reference RNA (Stratagene, Agilent Technologies Inc.) were labeled with Cy5 and Cy3, respectively, using the Low Input Quick Amp Labeling Kit (Agilent Technologies). Labeled cRNA was purified and quantified as described above. cRNA (300 ng per sample) and reference cRNA were hybridized to SurePrint G3 Mouse GE 8 x 60 K microarrays (Agilent Technologies) at 65 °C for 17 hours at 10 rpm.

Microarray analysis for FFPE tissues was performed according to our published one-color and two-color cDNA input protocols (Jackson et al. 2013). For the one-color FFPE protocol, 1650 ng of cDNA was labeled with 2.48 ul Cy3 dye using the Genomic DNA ULS Labeling Kit (Agilent Technologies). For the two-color FFPE protocol, 825 ng of cDNA was labeled with 1.44 ul Cy5 dye, and 825 ng of reference cDNA (made from a pool of all sample cDNA) was labeled with 1.24 μl Cy3 using the Genomic DNA ULS Labeling Kit (Agilent Technologies). In both protocols, excess dye was removed using KREApure columns (Agilent Technologies), labeled product was quantified using a NanoDrop spectrophotometer (Thermo Fisher Scientific Inc.), and degree of labeling was calculated using the following calculation: degree of labeling = [(340 × pmol per μl dye)/(ng per μl cDNA × 1000)] × 100% (with an acceptable range of 1.5-3%).

Hybridization mixtures were prepared using the Gene Expression Hybridization kit (Agilent Technologies) and CGHBlock (Agilent Technologies). For the one-color FFPE protocol, 600 ng of Cy3-labelled cDNA was hybridized to SurePrint G3 Mouse GE 8x60K microarrays (Agilent Technologies). For the two-color FFPE protocol, 300 ng each of sample and reference cDNA was hybridized to SurePrint G3 Mouse GE 8x60K microarrays (Agilent Technologies). Hybridization for both protocols occurred in a randomized block design, at 65°C for 17 hours at 20 rpm in the dark in an Agilent SureHyb hybridization chamber.

All slides were washed with Wash Buffers 1 and 2 (Agilent Technologies), and scanned at 5 μm resolution on an Agilent G2505B scanner. Feature extraction was accomplished using Agilent feature extraction software version 11. The complete microarray dataset is available through the Gene Expression Omnibus http://www.ncbi.nlm.nih.gov/geo/) accession number: GSE62843.

### Microarray statistical analysis

#### One-Color

Non-background subtracted median signal intensities were cyclic-LOWESS normalized (Bolstad et al. 2003) in R (R-Core-Development-Team 2012) using the affy library (RW.ERROR - Unable to find reference:449). Differential gene expression analysis between the control and treated samples was applied using the MAANOVA library (Wu et al. 2003) in R. The Fs statistic (Cui et al. 2005) was used to test for differential expression. The p-values from these tests were estimated using the permutation method with residual shuffling, and adjusted for multiple comparisons by using the false discovery rate (FDR) approach (Benjamini and Hochberg 1995). Least square means (Searle et al. 1980; Goodnight and Harvey 1978) were used to estimate the fold change for each pairwise comparison tested.

#### Two-color

A reference design (Kerr 2003) was employed for the two-color analysis. Non-background subtracted median signal intensities were LOWESS-normalized (Yang et al. 2002) in R (http://www.R-project.org/) using the MAANOVA library (RW.ERROR - Unable to find reference:449). The MAANOVA library was used for the DEG analysis using the log2 of the relative intensities. The p-values were estimated using the permutation method with residual shuffling, and FDR-adjusted p-values were computed using the FDR approach. Fold change estimates were computed as in the one-color analysis.

#### Correlation analysis

Correlations in figure 1 were determined using the regression function in the Microsoft Excel ‘Data Analysis’ tool pack.

### Microarray bioinformatic analysis

#### Cluster analysis

The universal reference used in the two-color FRO data was different than the pooled sample reference used in the FFPE two-color analyses. For the cluster analysis, the data from the two-color and one-color experiments were normalized to the median of the time-matched controls. By applying this normalization the FRO and FFPE data could then be analysed jointly. Heatmaps were constructed using a similarity metric based on 1-correlation. The correlations were estimated using the Spearman approach. Genes were considered significant with a 2-fold change and an FDR p < 0.05. The row dendrogram for the heatmaps was estimated by pooling the one-color and two-color data in order to have the same row order for the two plots to help facilitate the comparison of the two heatmaps.

#### Pathway analysis

Ingenuity Pathway Analysis (IPA; http://www.ingenuity.com/) was used to identify enriched molecular pathways. Gene expression data was analyzed using an IPA Core Analysis with a gene expression threshold of fold change ≥ ±1.5 and FDR p ≤ 0.05. Enrichment of IPA canonical pathways was determined based on the number of DEGs in the dataset that also appear in that pathway. Pathway significance thresholds were determined in IPA using a right-tailed Fisher’s exact test. Only mapped DEGs were considered for figure 2 (unmapped DEGs in Suppl. Fig. S1).

#### NextBio

To identify chemical effectors and disease states that produce changes to gene expression that are similar to those produced following exposure to furan, gene expression profiles from all microarray data sets were mined against a large genomic database repositories using NextBio (www.nextbio.com). NextBio uses proprietary algorithms that use pairwise gene signature correlations and rank-based enrichment statistics to produce scores, where the chemical or disease with the highest similarity is given a score of 100 and the rest are normalized accordingly (Kupershmidt et al. 2010). The meta-analyses included up to the top 250 DEGs (mapped) from each group with an FDR p < 0.05.

### RNA-sequencing methods

RNA sequencing was performed on the FRO and FFPE tissues using two different protocols: ribosomal RNA-depleted-sequencing (rmRNA-seq; conducted in the Mason laboratory, NYC, USA) and poly-A-selected RNA-sequencing (mRNA-seq; conducted at Genome Quebec, Montreal, Canada). The rmRNA-seq and mRNA-seq protocols were carried out according to Illumina’s TruSeq Total RNA Sample Prep Kit and Illumina’s TruSeq RNA Sample Preparation Kit v2, respectively. For rmRNA-seq library construction, ribosomal RNA was removed using biotinylated, target-specific oligos combined with Ribo-Zero rRNA removal beads (Human/Mouse/Rat). For mRNA-seq library construction, poly-A-containing mRNA was purified using oligo-dT-attached magnetic beads. Following purification, rRNA-depleted or poly-A-enriched RNA was chemically fragmented into small pieces using divalent cations under elevated temperature. For all FFPE samples, we used double the input when compared to fresh-frozen. The RNA fragments were copied into first strand cDNA using reverse transcriptase and random hexamer primers, followed by second strand cDNA synthesis using DNA Polymerase I and RNase H. Adapters were subsequently ligated to the cDNA fragments, and then purified and enriched for with PCR to create a final cDNA library. The final cDNA libraries were single-end sequenced on a HiSeq2000. Ribo-depletion experiments had a read length of 42 bases and a read depth of 10-60M reads. Poly-A-enrichment experiments had a read length of 50 bases and a read depth of 3.8-11M reads. It has previously been reported that the required sequencing depth to be microarray-equivalent is 5M (Black et al. 2014). The complete dataset is available through the Gene Expression Omnibus http://www.ncbi.nlm.nih.gov/geo/) accession number: GSE62843.

### RNA-seq bioinformatic analysis

The RNA-sequencing data were processed using Illumina’s RTA software and converted into FASTQ files using Illumina’s CASAVA pipeline. The FASTQ files were aligned to either the UCSC mm9 mouse genome (for the furan study) or the rn5 rat genome (for the age-in-block study) with STAR (Dobin et al. 2013), a universal RNA-seq aligner, using default parameters. Sequences that mapped to more than one locus were excluded from further analysis. Gene expression values were calculated based on composite gene models of NCBI’s Reference Sequence (RefSeq) gene annotation models (retrieved via UCSC’s Table Browser 5/7/14) using featureCounts (Liao et al. 2014), a program for assigning sequence reads to genomic features. Composite gene models for each gene consisted of the union of the exons from the set of all transcript isoforms from that gene. Expression values were also calculated for a subset of RefSeq genes, which corresponded to the intersection between the RefSeq gene models and the Agilent SurePrint G3 Mouse Microarray targets. The intersection between the RefSeq gene models and the Agilent SurePrint G3 Mouse Microarray targets was determined as follows. Microarray probe sequences corresponding to Agilent SurePrint G3 Mouse Microarrays were downloaded from NCBI (accession # GPL13912). The probe sequences were aligned to the mouse genome using STAR. Probe sequences that mapped to more than one locus (5.65%) were excluded from downstream consideration. Probe sequences that mapped to just one locus were intersected with composite gene models from NCBI’s RefSeq database using featureCounts. Probe sequences that unambiguously overlapped with no more than one RefSeq composite gene model (53.9%) were paired with that gene model; the remaining probe sequences were discarded. The pairs were collated into a unique, non-redundant list of RefSeq genes, which was used in subsequent differential gene expression analyses. All sets of gene counts were normalized using voom (Law et al. 2014), which performs a LOWESS regression to estimate the mean-variance relation and transforms the gene counts into the appropriate log form for linear modeling. Differential gene expression analysis was then performed for each set of transformed gene counts using limma (Ritchie et al. 2015) for the following set of contrasts: FF Control vs. FF High Dose, FFPE Control 18hrs vs. FFPE High Dose 18hrs, and FFPE Control 3wks vs. FFPE High Dose 3wks.

#### Online data access

The complete dataset is available through the Gene Expression Omnibus (www.ncbi.nlm.nih.gov/geo/). The superseries accession number is GSE62843.

#### Disclosure declaration

The authors declare no conflicts of interest. The research described in this article has been reviewed by the U.S. EPA and approved for publication. Approval does not signify that the contents necessarily reflect the views or the policies of the Agency. Mention of trade names or commercial products does not constitute endorsement or recommendation for use.

## Acknowledgements

We thank the ILS animal care group for performing the furan mouse exposures and fixing these tissues, and the NTP contract NO1-ES-55536 for preparing the archival RNA. This research was funded by HESI, the Federal Government of Canada’s Genomics R&D Initiative, the Canadian Regulatory Systems for Biotechnology, and the U.S. EPA Office of Research and Development. A.F.W. was supported by the Natural Science and Engineering Research Council of Canada and the Ontario Graduate Scholarship. This work was supported with funding from the National Institutes of Health (NIH), including R01NS076465, as well as funds from the Irma T. Hirschl and Monique Weill-Caulier Charitable Trusts, the STARR Consortium (I7-A765). The authors greatly acknowledge Weill Cornell Epigenomics Core contribution and technical support from Jennifer A. Busuttil and Caroline Sheridan. We also thank the Bert L. and N. Kuggie Vallee Foundation Young Investigator Award and the WorldQuant Foundation. The authors greatly acknowledge Dr. Matt Meier and Dr. Ivy Moffat for helpful comments on the manuscript.

## Figure Legends

**Suppl. Fig. S1.**
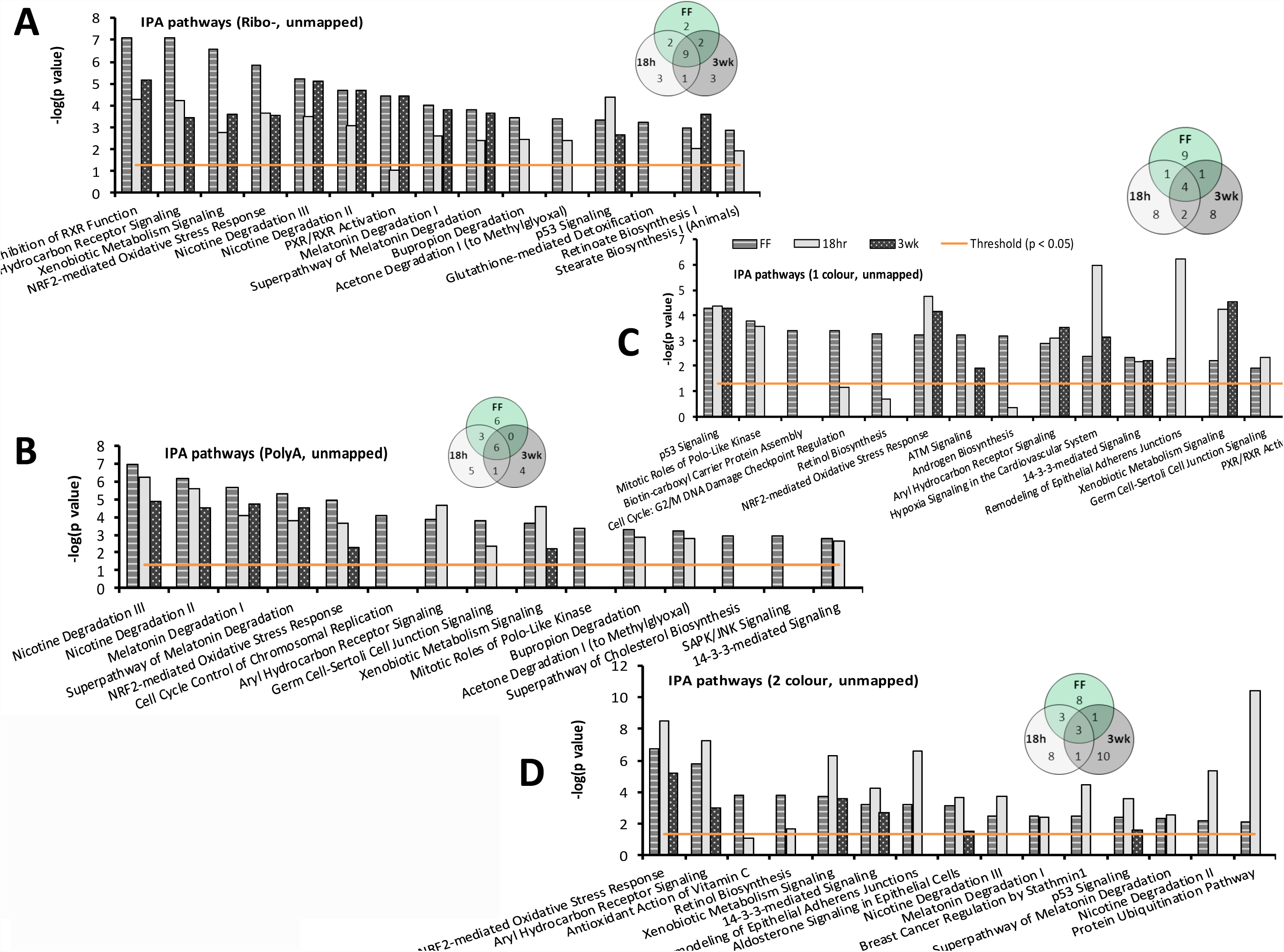
Pathway analysis for (A) RNA-seq ribo-depletion, (B) RNA-seq poly-A-enrichment, (C) one-color microarrays, and (D) two-color microarrays, showing the top 15 enriched pathways for the fresh-frozen (FRO) samples and enrichment levels of these pathways in the corresponding FFPE samples. Pathway significance was calculated in IPA using a Fisher’s Exact test (-log(p-value)=1.3 corresponds to a p=0.05). Venn diagrams show the overlap of the top 15 pathways in each fresh-frozen, FFPE-18hr, and FFPE-3wk. For each analysis, all DEGs (including unmapped) with FDR p < 0.05 and fold change > +/-1.5 were used.

---

**Supplemental Table S1.** RNA quality.

| **Time in formalin** | **Animal ID** | **[RNA]**<br>**(ng/ul)** | **260/280** | **260/230** | **RIN** |
| --- | --- | --- | --- | --- | --- |
| Fresh-frozen | 2 | 150.75 | 2.07 | 2.24 | 9.5 |
| Fresh-frozen | 3 | 153.06 | 2.08 | 2.25 | 9.3 |
| Fresh-frozen | 4 | 163.21 | 2.07 | 2.23 | 9 |
| Fresh-frozen | 6 | 153.88 | 2.05 | 2.19 | 9.3 |
| Fresh-frozen | 41 | 157.81 | 2.02 | 2.27 | 9.5 |
| Fresh-frozen | 42 | 145.53 | 2.06 | 2.16 | 9.2 |
| Fresh-frozen | 43 | 155.736 | 2.05 | 2.25 | 9.4 |
| Fresh-frozen | 44 | 148.05 | 2.03 | 2.06 | 8.9 |
| 18 hours | 2 | 253.55 | 2.06 | 2 | 2.1 |
| 18 hours | 3 | 315.79 | 2.06 | 1.92 | 2 |
| 18 hours | 4 | 173 | 2.12 | 2.05 | 2.2 |
| 18 hours | 6 | 318.29 | 2.05 | 1.99 | 2 |
| 18 hours | 41 | 275.76 | 2.05 | 1.93 | 2 |
| 18 hours | 42 | 429.72 | 2.06 | 1.97 | 2 |
| 18 hours | 43 | 276.67 | 2.08 | 2.06 | 1.9 |
| 18 hours | 44 | 300.55 | 2.06 | 2 | 2.3 |
| 3 weeks | 2 | 36.82 | 1.79 | 1.33 | 2.4 |
| 3 weeks | 3 | 43.14 | 1.91 | 1.84 | 2.4 |
| 3 weeks | 4 | 36.89 | 1.81 | 2.11 | 2.4 |
| 3 weeks | 6 | 31.87 | 1.78 | 1.94 | 2.4 |
| 3 weeks | 41 | 27.11 | 1.85 | 1.88 | 1.3 |
| 3 weeks | 42 | 17.28 | 1.89 | 1.88 | 2.2 |
| 3 weeks | 43 | 8.8 | 1.74 | 2.32 | 2.15 |
| 3 weeks | 44 | 27.48 | 1.87 | 2.03 | NA |

